# Ligandability Assessment of the LAG-3 D1 Domain Enables Discovery of a Small-Molecule Inhibitor

**DOI:** 10.64898/2025.12.07.692800

**Authors:** Nelson García-Vázquez, Moustafa T. Gabr

## Abstract

LAG-3 is an emerging immune checkpoint whose extracellular D1 domain engages MHC class II through a broad protein-protein interface traditionally considered difficult to modulate with small molecules. To evaluate the ligandability of this region, we combined 100-ns molecular dynamics (MD) simulations, structure-based virtual screening, and biophysical and biochemical assays. MD sampling of the isolated D1 domain revealed a recurrent, transient cavity adjacent to the MHCII-binding surface. A representative pocket-open conformation was used to screen a ∼10,240-compound diversity library, yielding a single validated hit, N05. N05 bound the D1 domain with micromolar affinity measured by microscale thermophoresis (Kd = 59.2 µM, TRIC/MST channel) and by spectral-shift detection (Kd = 56.1 µM), and it partially inhibited the LAG-3/MHCII interaction (EC_50_ = 42.9 µM; maximal inhibition ∼76%). A 30-ns MD simulation of the LAG-3-N05 complex showed stable ligand engagement within the MD-identified cavity and consistent stabilizing interactions with residues forming the pocket. These results demonstrate that the LAG-3 D1 domain possesses an accessible, dynamically formed binding site capable of accommodating small molecules, providing a structural and biophysical foundation for future exploration of LAG-3 ligandability.

**Graphical Abstract:** 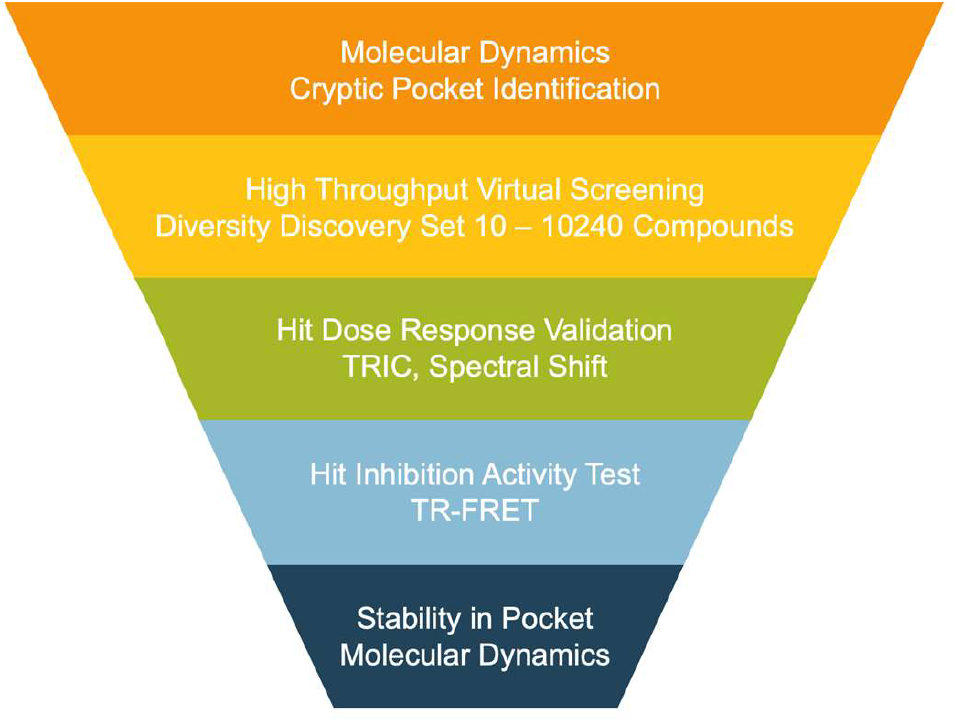

Overview of the computational and experimental workflow used to identify and validate a small-molecule binder of the LAG-3 D1 domain.

## A. Introduction

Immune checkpoint blockade has transformed the treatment of multiple cancers, primarily through monoclonal antibodies (mAbs) targeting PD-1, PD-L1 and CTLA-4. However, a substantial fraction of patients fail to respond or develop resistance, and antibody therapies are associated with high cost, parenteral administration, and immune-related adverse events. These limitations have motivated efforts to expand the repertoire of targeted pathways and to explore alternative therapeutic modalities, including small molecules, peptides, and degraders that modulate immune checkpoints through mechanisms distinct from classical antibody blockade.^1-4^

Lymphocyte activation gene-3 (LAG-3, CD223) is an inhibitory receptor expressed on activated CD4^+^ and CD8^+^ T cells, regulatory T cells, and other lymphocyte subsets, where it acts as a key regulator of T-cell exhaustion and tolerance in cancer and autoimmunity.^1,2,5-7^ The extracellular region of LAG-3 contains four Ig-like domains (D1-D4); domain 1 (D1) is unusual in harboring a proline-rich loop that mediates high-affinity binding to MHC class II (MHCII).^1,5^ Recent structural and functional studies have clarified how LAG-3 engages MHCII and additional ligands such as fibrinogen-like protein-1 (FGL1), revealing that LAG-3 dampens T-cell activation by competing with CD4 for MHCII binding, altering immunological synapse geometry, and transmitting inhibitory signals through its cytoplasmic tail.^5-9^ Clinically, LAG-3-directed mAbs and bispecifics combined with PD-1 blockade have shown encouraging activity in multiple tumor types, underscoring the therapeutic relevance of this pathway.^2,3^

High-resolution structural information has now become available for the human LAG-3 ectodomain, for LAG-3 in complex with MHCII, and for LAG-3 bound to therapeutic antibodies.^5-7^ These studies show that D1 directly contacts a conserved, membrane-proximal region of MHCII spanning the α2 and β2 subdomains and that LAG-3 dimerization constrains MHCII spacing at the cell surface.^5,7^ Antagonist antibodies that bind the flexible loop in D1 can simultaneously block MHCII and FGL1 engagement and potently relieve LAG-3-mediated inhibition.^5,9^ Together, these data strongly implicate the D1 region, and in particular the MHCII-engaging surface, as a functionally privileged site for therapeutic intervention.

Despite this progress, the ligandability of LAG-3 by small molecules remains poorly characterized. Most clinically advanced LAG-3 inhibitors are antibodies or protein-based agents,^2,3^ whereas small-molecule checkpoint programs have so far focused largely on PD-1/PD-L1, where multiple chemotypes identified by structure-based design, DNA-encoded libraries, and pharmacophore methods can disrupt the receptor-ligand interaction and restore T-cell activity.^4,12-14^ Even in those systems, early hits often exhibit micromolar affinity yet provide crucial proof of druggability and starting points for optimization.^12-14^ For LAG-3, the available crystal and cryo-EM structures describe an extended, solvent-exposed protein-protein interface between D1 and MHCII with few obvious pockets in static views,^5-7^ which has contributed to the perception that this surface may be intrinsically challenging for conventional small-molecule binding.

Over the past decade, molecular dynamics (MD) simulations have emerged as a powerful approach to uncover “cryptic” or transient binding pockets that are not apparent in static structures but become accessible as proteins explore their conformational landscape.^15-18^ MD and enhanced-sampling methods, sometimes combined with mixed-solvent or fragment probes, have revealed druggable cryptic sites in diverse targets previously considered undruggable, and in several cases these pockets have been validated by fragment screening and crystallography.^15,17,18^ Recent AI-driven tools such as PocketMiner further suggest that cryptic pockets may be widespread across the human proteome, substantially expanding the set of potentially ligandable proteins.^16^ These conceptual advances raise the possibility that LAG-3 D1 may transiently form pockets adjacent to, or overlapping with, the MHCII-binding surface that could be exploited by appropriately designed small molecules.

Here, we set out to ask whether the D1 domain of LAG-3 harbors a ligandable pocket that can be targeted by small molecules, using an integrated computational-to-biophysical workflow. Guided by MD simulations of the LAG-3 D1 region, we identified a transient pocket in the vicinity of the MHCII-binding surface and subjected this site to structure-based virtual screening of a focused ∼1,000-member library. Top-ranked candidates were evaluated in a biochemical/biophysical assay, leading to the identification of a micromolar-affinity binder that partially disrupts LAG-3 function in vitro. Follow-up MD simulations were then used to probe the stability of the ligand-pocket interaction and to map key residue contacts over time. Rather than aiming to deliver a fully optimized inhibitor, this work is intended as a feasibility study that (i) assesses the small-molecule ligandability of LAG-3 D1, (ii) illustrates how MD-derived pockets can be coupled to virtual screening for a challenging immune checkpoint target, and (iii) provides an initial structural framework and tractable hit series for future medicinal chemistry and mechanistic studies.

## B.Materials and methods

### Virtual Screening

A receptor structure was prepared by first selecting a representative pocket-open conformation consistent with the cryptic cavity identified in MD sampling. The protein model was prepared using the Schrödinger Protein Preparation Wizard: hydrogens were added, protonation states optimized at pH 7.4 ± 0.2, and restrained minimization was applied. The docking grid was centered on the residues defining the cryptic pocket region.

The Enamine Discovery Diversity Set (10,240 compounds, SDF format) was processed using LigPrep with Epik to generate low-energy protonation states around pH 7.4 ± 2.0. Docking was performed using Schrödinger Glide with a three-stage funnel: HTVS → SP → XP. At each stage, the top 10% of scored compounds were advanced, resulting in nine final XP-ranked candidates.

### Microscale Thermophoresis / Spectral Shift Binding Assay

Compound N05 was obtained from Enamine (catalog code Z3071585108) as a powder and dissolved in DMSO to prepare a 50 mM stock solution. For binding assays, the stock was diluted such that the final DMSO concentration never exceeded 3% (v/v).

Recombinant human LAG-3-His was purchased (SinoBiological, Cat. 16498-H08H) and labeled using the RED-tris-NTA 2nd Generation dye (NanoTemper MO-L018), following the manufacturer’s instructions. Labeled protein (50 nM final) was diluted in PBS (pH 7.4) with 0.05% Tween-20. N05 was titrated in a 12-point half-dilution series starting at 200 µM. After 15 min incubation at room temperature, thermophoresis and spectral-shift signals were recorded on a NanoTemper Monolith X using default acquisition settings. Data were analyzed with the vendor’s software to extract apparent dissociation constants.

### HTRF LAG-3 / MHCII Inhibition Assay

Functional inhibition of the LAG-3-MHCII interaction was measured using the Revvity HTRF Human LAG-3 / MHCII Binding Kit per manufacturer instructions. N05 was tested in a 12-point half-dilution series beginning at 200 µM. After mixing assay components, plates were incubated for 3 hrs at room temperature before fluorescence detection. Four-parameter logistic fitting was used to determine EC_50_ values and maximal inhibition.

#### Protein Preparation for MD

The structural model for molecular dynamics simulations was derived from PDB 9CYM, which contains the extracellular LAG-3 D1-D2 module in complex with MHCII. Only the LAG-3 polypeptide was retained, and the construct was truncated to include only the D1 domain.

Missing residues in the 69-92 loop were built using CHARMM-GUI; all native disulfide bonds in D1 were preserved. The prepared monomeric D1 domain served as the basis for apo MD simulations.

For complex simulations, the docking-derived pose of N05 was used; ligand parameters were generated using CHARMM-GUI.

#### Molecular Dynamics Simulations

All simulations were carried out using GROMACS 2025.3 with the CHARMM36m force field. System preparation (topology, solvation, ion placement) was done via CHARMM-GUI. Protein or protein-ligand systems were solvated in an explicit TIP3P water box with a 10 Å padding and neutralized to 150 mM NaCl using distance-based ion placement.

After energy minimization, systems underwent equilibration under NVT and NPT ensembles following CHARMM-GUI defaults. Temperature was maintained at 300 K using the V-rescale thermostat, and pressure coupling (during NPT) used the barostat defined in the CHARMM-GUI equilibration scripts. Covalent bonds involving hydrogens were constrained via LINCS, and long-range electrostatics were treated with PME. A 2-fs timestep was used for production runs under the Verlet cutoff scheme.

Simulations comprised a 100-ns apo run for the isolated D1 domain and a 30-ns run for the LAG-3-N05 complex, both under identical conditions.

#### MM-GBSA Binding Energy and Per-Residue Decomposition

To estimate binding free energy and per-residue contributions, we used gmx_MMPBSA on frames sampled from the production trajectory. Standard GB parameters compatible with CHARMM36m were applied; both total binding free energies and residue-wise decompositions were computed over the same subset of frames.

## C. Results

### 1. MD Analysis Reveals a Transient Cryptic Pocket in the LAG-3 D1 Domain

To evaluate whether the D1 domain of LAG-3 harbors cryptic structural features capable of supporting small-molecule binding, we performed a 100-ns all-atom MD simulation of the isolated D1 module, starting from the crystal structure of the human LAG-3 ectodomain (PDB 9CYM). The structure was prepared in CHARMM-GUI by extracting a monomeric D1 unit, adding missing loop residues 69-92, introducing native disulfide bonds, solvating the protein in a rectangular TIP3P water box with 10 Å padding, and neutralizing the system with 150 mM NaCl. Production MD was run in GROMACS 2025.3 using standard CHARMM36m parameters following energy minimization and NVT/NPT equilibration (see Methods).

A visual overview of the D1 domain and the MD-identified cavity is shown in **Fig. 1A-C**.

**Figure 1.**
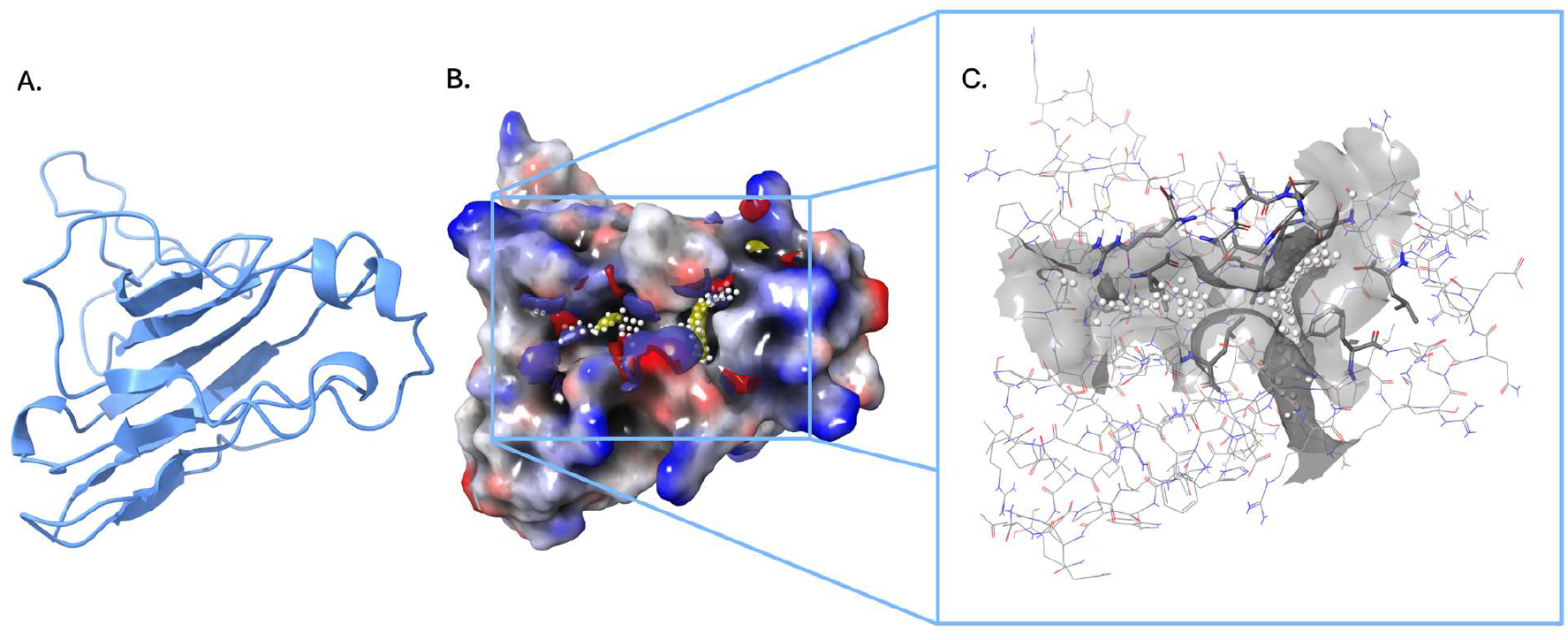
Identification of a cryptic pocket on the LAG-3 D1 domain through molecular dynamics simulation. **(A)** Ribbon representation of the LAG-3 D1 domain used as the initial structure for MD simulations. **(B)** Electrostatic surface map of D1 highlighting the transient pocket observed during MD sampling (yellow spheres indicate pocket-lining atoms). **(C)** Enlarged view of the MD-identified cavity showing the pocket geometry and surrounding residues rendered in surface and stick representation.

Over the course of the apo simulation, we observed reproducible opening of a shallow but clearly defined cavity on the face of D1 that interacts with MHCII (**Fig. 1B**). The cavity appeared intermittently throughout the trajectory, opening and closing in a manner characteristic of dynamic “cryptic” pockets rather than a constitutively present binding site. Pocket formation occurred at the interface of several flexible loops and β-strand edges that contribute to MHCII recognition. Notably, the cavity formed adjacent to, but not directly within, the proline-rich loop responsible for high-affinity MHCII binding, suggesting that small molecules engaging the transient cavity could allosterically modulate the MHCII-binding surface without directly competing with the conserved pMHCII interaction.

Inspection of frames in which the cavity was open revealed a recurring set of residues that form its walls and floor (**Fig. 1C**), including CYS44, SER45, and PRO46 in the N-terminal region; ALA59, GLY60, VAL61, THR62, SER54, SER55, and ARG57 in the central groove; PHE132, SER133, and LEU134 forming a hydrophobic-aromatic belt; and ALA149, ALA150, VAL151, LEU158, SER159, and CYS160 along the C-terminal flank. Additional surrounding residues, ASP131, GLU124, ARG127, and GLY126, further shaped the cavity. Together, these residues create a mixed hydrophobic-polar environment consistent with a partially solvent-shielded site capable of accommodating small heterocycles or fragment-sized ligands. Although the cavity remained relatively shallow, it reformed multiple times with comparable geometry across the trajectory, indicating a structurally permissive region capable of supporting ligand binding.

The MD-revealed cavity emerges at, or immediately adjacent to, the functional MHCII-facing surface of D1, a region traditionally regarded as a difficult, solvent-exposed protein-protein interface. The repeated spontaneous formation of this pocket suggests latent ligandability and supports the idea that LAG-3 D1 contains cryptic structural features amenable to small-molecule intervention. A representative open-pocket conformation consistent with this MD-identified cavity was therefore selected as the receptor for downstream virtual screening (see Section 2 and **Fig. 1B-C**).

### 2. Virtual Screening of the MD-Revealed Pocket Identifies a Single Ligand (N05)

To evaluate whether the transient cavity observed in the MD simulations could accommodate small-molecule binders, we performed a structure-based virtual screen centered on this site. A representative pocket-open conformation consistent with the MD-identified geometry was selected as the receptor, and the structure was prepared in Maestro using the Protein Preparation Wizard with standard settings, including hydrogen addition, optimization of protonation states at pH 7.4 ± 0.2, and restrained minimization. A Glide docking grid was then generated by selecting residues lining the cavity and, in parallel, by centering the grid on the ligand in the internal MD-derived reference conformation; both approaches produced effectively identical grid placement over the targeted region (**Fig. 2A**).

**Figure 2.**
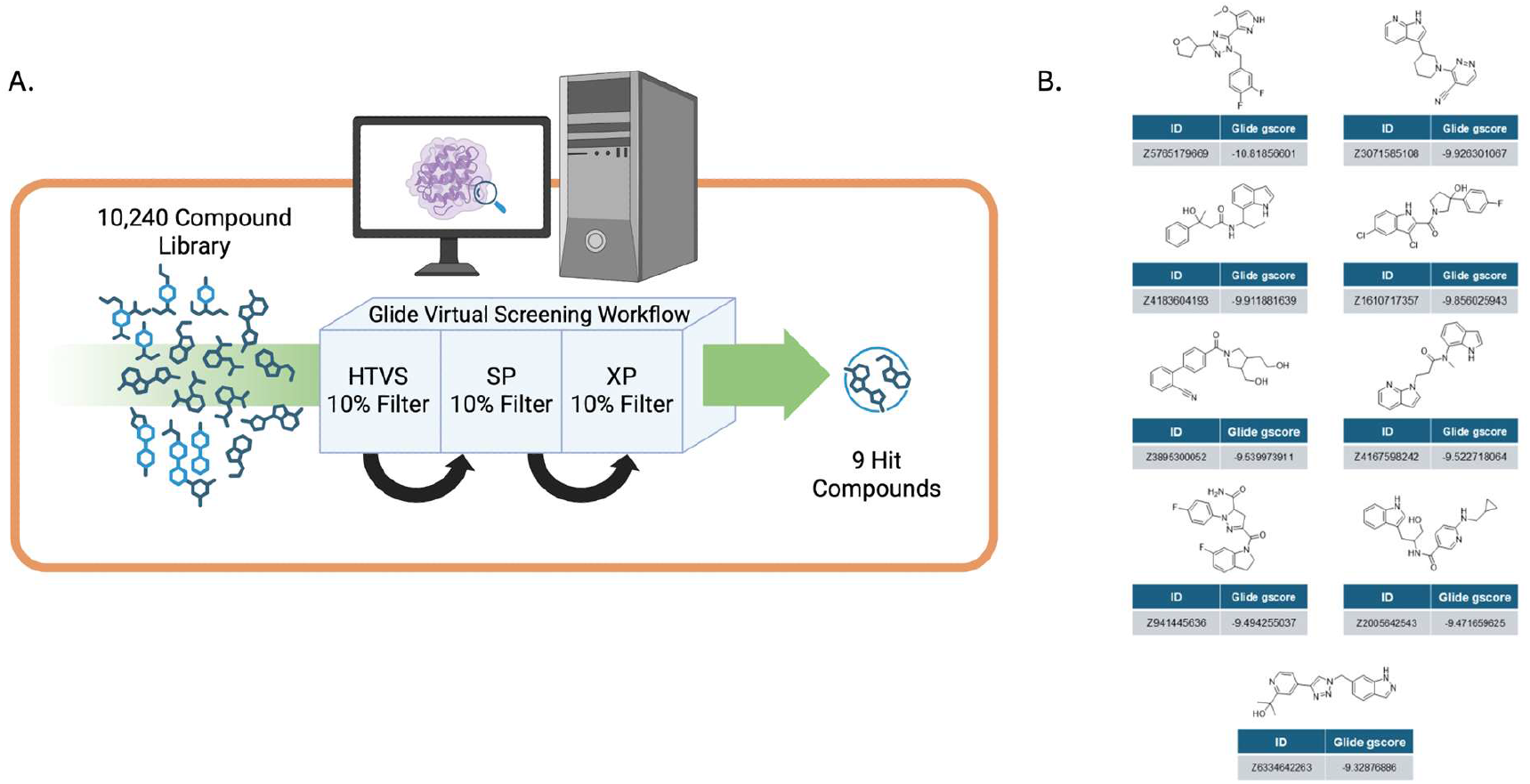
Virtual screening of the LAG-3 D1 pocket and resulting hits. **(A)** Schematic overview of the Glide virtual screening workflow. A 10,240-compound library was processed through sequential HTVS, SP, and XP docking stages, retaining the top 10% of compounds at each step. **(B)** Chemical structures and Glide XP scores of the nine top-ranked hit compounds obtained from the virtual screening funnel.

The Enamine DDS Library (10,240 compounds) was processed in LigPrep to generate 3D structures and relevant protonation states (Epik, pH 7.0 ± 2.0), after which compounds were docked through a three-stage funnel consisting of high-throughput virtual screening (HTVS), standard-precision (SP) docking, and extra-precision (XP) refinement. At each stage the top 10% of compounds were advanced, ultimately yielding nine XP-ranked candidates (**Fig. 2B**). These nine molecules were selected strictly according to their XP scores, without manual rescoring, clustering, or pharmacophore-based filtering.

### 3. Biophysical Assays Confirm Micromolar Binding of N05 to LAG-3 D1

To assess the computationally selected hits, all shortlisted compounds were evaluated by microscale thermophoresis (MST) and TRIC measurements using recombinant human LAG-3-His labeled with RED-tris-NTA dye. Each compound was titrated in a 12-point half-dilution series beginning at 200 µM, providing a concentration range suitable for detecting low- to high-micromolar binding events. Across these assays, the thermophoretic and spectral-shift responses were examined for clear, concentration-dependent behavior while monitoring for potential artifacts such as aggregation, nonspecific interference, or solubility limitations, none of which were observed under these conditions.

Among the nine candidates tested, only Z3071585108, designated N05, produced reproducible dose-dependent response curves in both detection channels (**Fig. 3A-B**) with Kd of 59.2 µM (TRIC) and 56.1 µM (spectral shift). Based on this consistent and quantifiable binding behavior, N05 was selected for subsequent functional inhibition assays.

**Figure 3.**
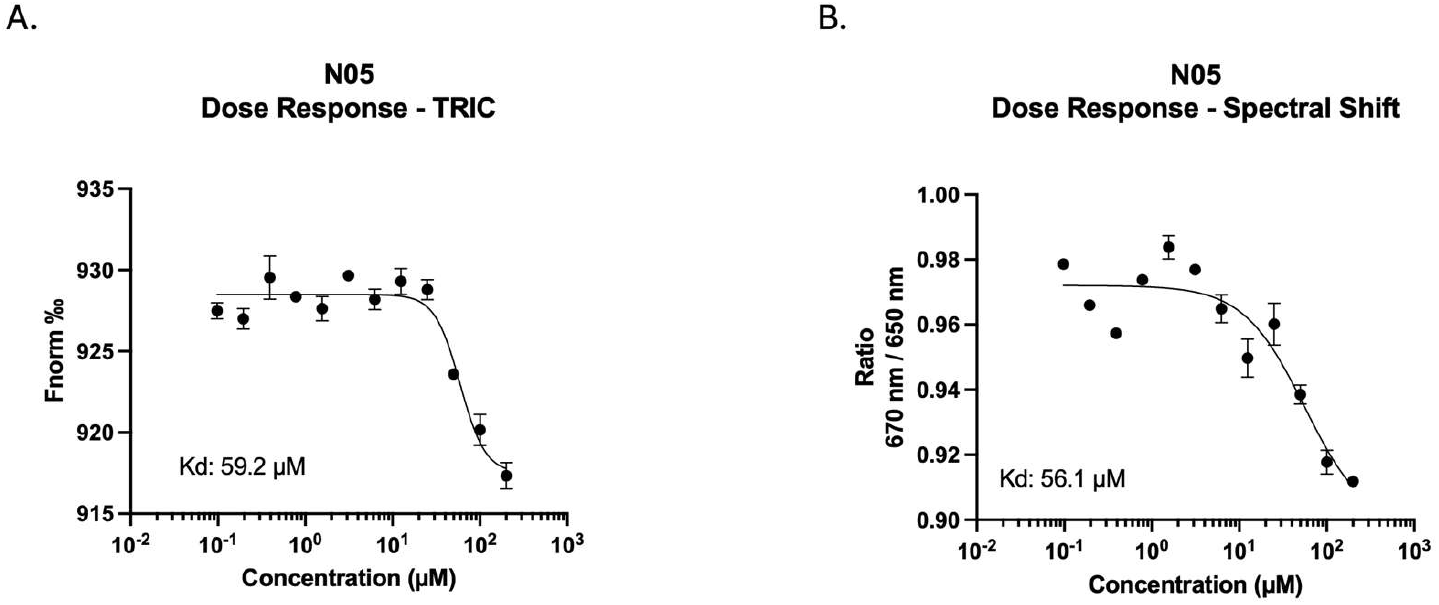
Biophysical evaluation of hit compound N05. **(A)** MST TRIC dose response curve for N05 binding to LAG-3 D1, yielding an apparent Kd of 59.2 µM. **(B)** Spectral-shift dose response curve measured under the same conditions, yielding an apparent Kd of 56.1 µM.

### 4. N05 Partially Inhibits the LAG-3/MHCII Interaction in an HTRF Binding Assay

To assess whether N05 binding to the D1 domain translates into functional modulation of LAG-3, the compound was tested using a time-resolved FRET-based HTRF assay that measures disruption of the LAG-3-MHC class II interaction (**Fig. 4A**). Experiments were performed according to the kit protocol, with compound preincubation under standard assay conditions.

**Figure 4.**
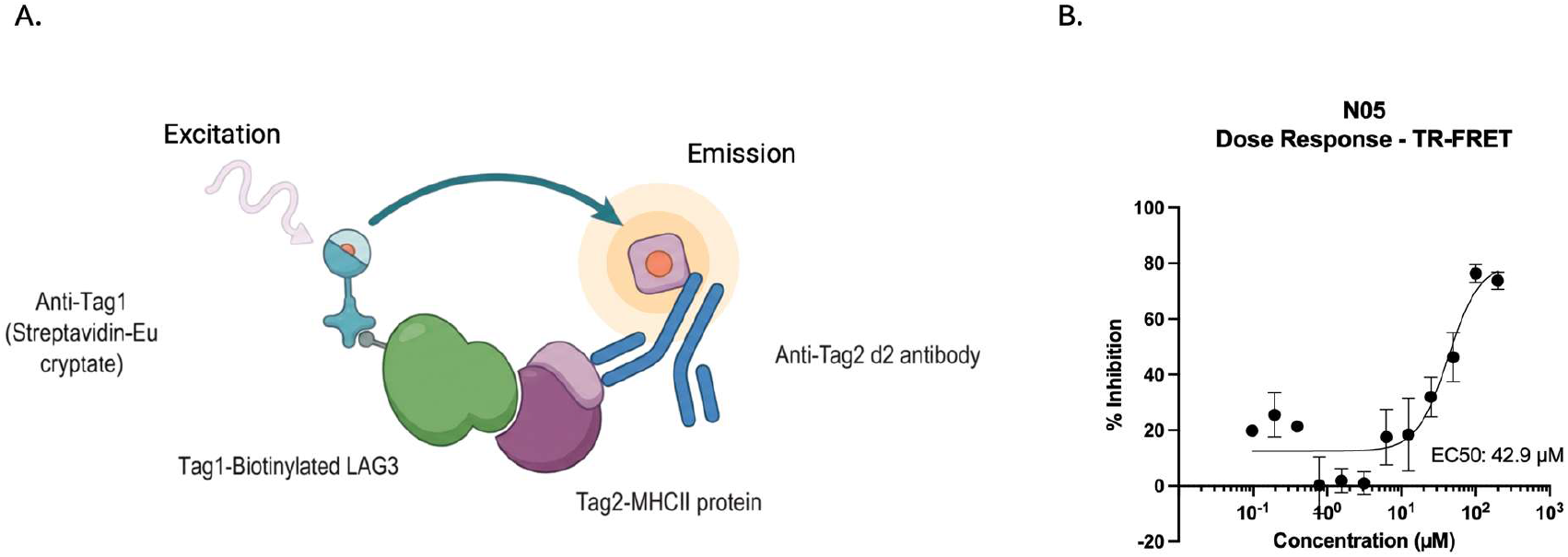
N05 partially inhibits the LAG-3-MHCII interaction. **(A)** Schematic overview of the TR-FRET assay used to measure disruption of the LAG-3-MHCII interaction. **(B)** Dose-response curve showing partial inhibition of LAG-3-MHCII binding by N05, with an EC_50_ of 42.9 µM and a maximal inhibition of ∼76%.

N05 elicited a clear, dose-dependent response consistent with inhibition of LAG-3 binding to MHCII (**Fig. 4B**) with an EC_50_ of 42.9 µM, and maximal inhibition of ∼76% at 100 µM. The partial plateau suggests that N05 does not fully abrogate the interaction under the conditions tested but is capable of substantially impairing LAG-3-MHCII interaction at higher concentrations.

### 5. MD Simulation of the N05-LAG-3 Complex Reveals Early Pose Adjustment Followed by Stable Pocket Engagement

To examine the conformational stability of N05 within the MD-identified pocket, we conducted a 30-ns all-atom MD simulation of the LAG-3 D1-N05 complex (**Fig. 5A-E**). The starting structure was taken from the induced-fit docking pose, and the system was parameterized using CHARMM-GUI with conditions matched to those used in the biophysical assays. After solvation and ion setup, the complex was simulated in GROMACS under CHARMM36m using the same equilibration and production workflow applied to the apo system.

**Figure 5.**
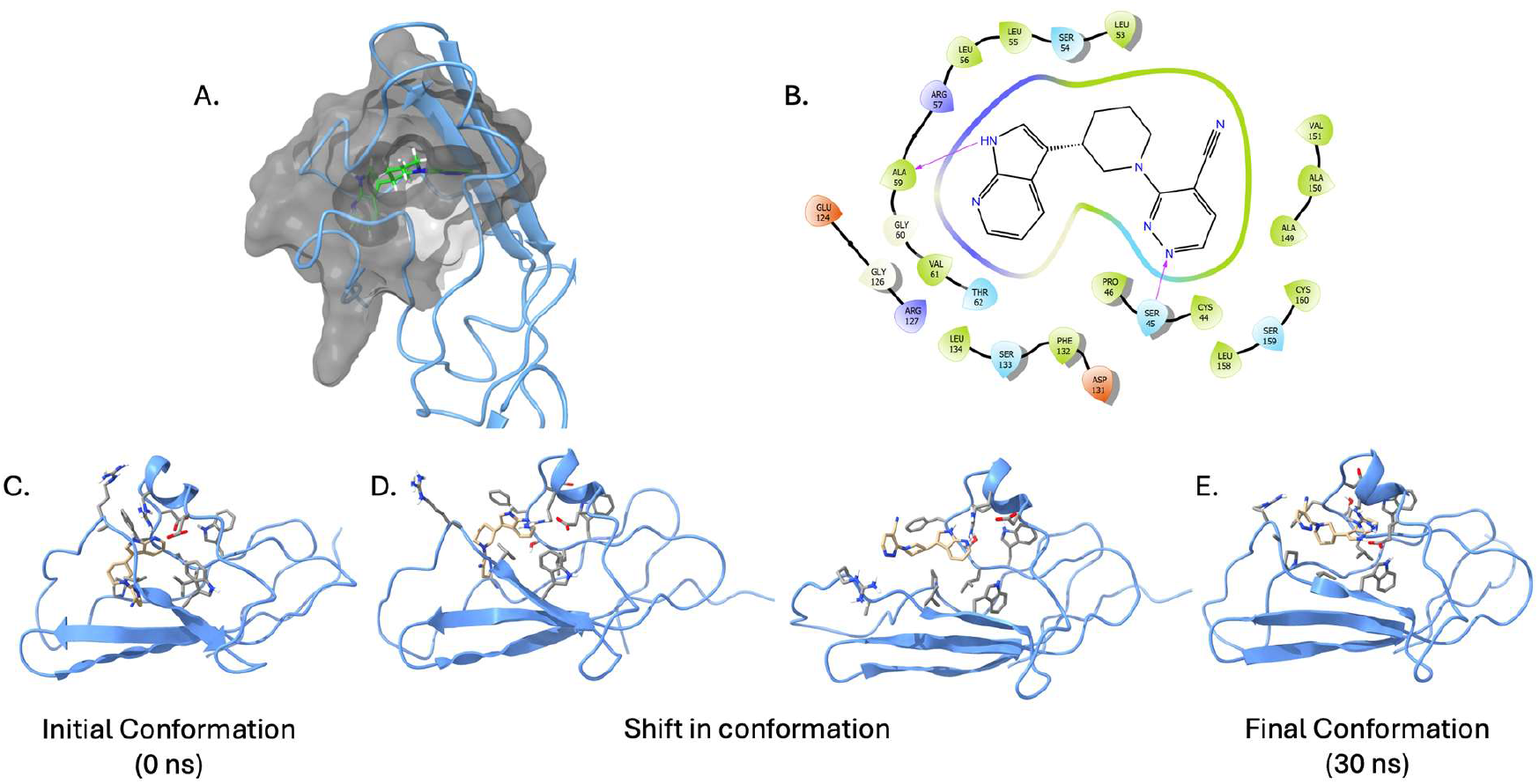
MD simulation of N05 in the LAG-3 D1 pocket reveals conformational adjustment and stable binding. **(A)** Surface representation of the MD-identified pocket with N05 bound in the refined pose. **(B)** Two-dimensional interaction map showing residues contacting N05. **(C)** Initial ligand conformation at the start of the MD trajectory (0 ns). **(D)** Intermediate frames illustrating ligand rearrangement within the pocket. **(E)** Final ligand conformation after 30 ns, showing stable engagement of the cryptic cavity.

Analysis of ligand RMSD (protein-fitted) showed an initial increase during the first 8-10 ns (**Fig. 6A**), reaching approximately 0.55-0.65 nm, which reflects early pose relaxation and conformational adjustment within the cavity. Following this phase, RMSD values stabilized for the remainder of the trajectory, indicating that the ligand adopted a consistent and persistent orientation in the pocket. A complementary ligand-pocket distance metric displayed a similar pattern (**Fig. 6B**), with an early transient increase to roughly 1.3-1.4 nm followed by relaxation and stabilization near 0.9-1.0 nm for most of the simulation. Over the final ∼20 ns, neither metric showed evidence of ligand dissociation or progressive destabilization.

**Figure 6.**
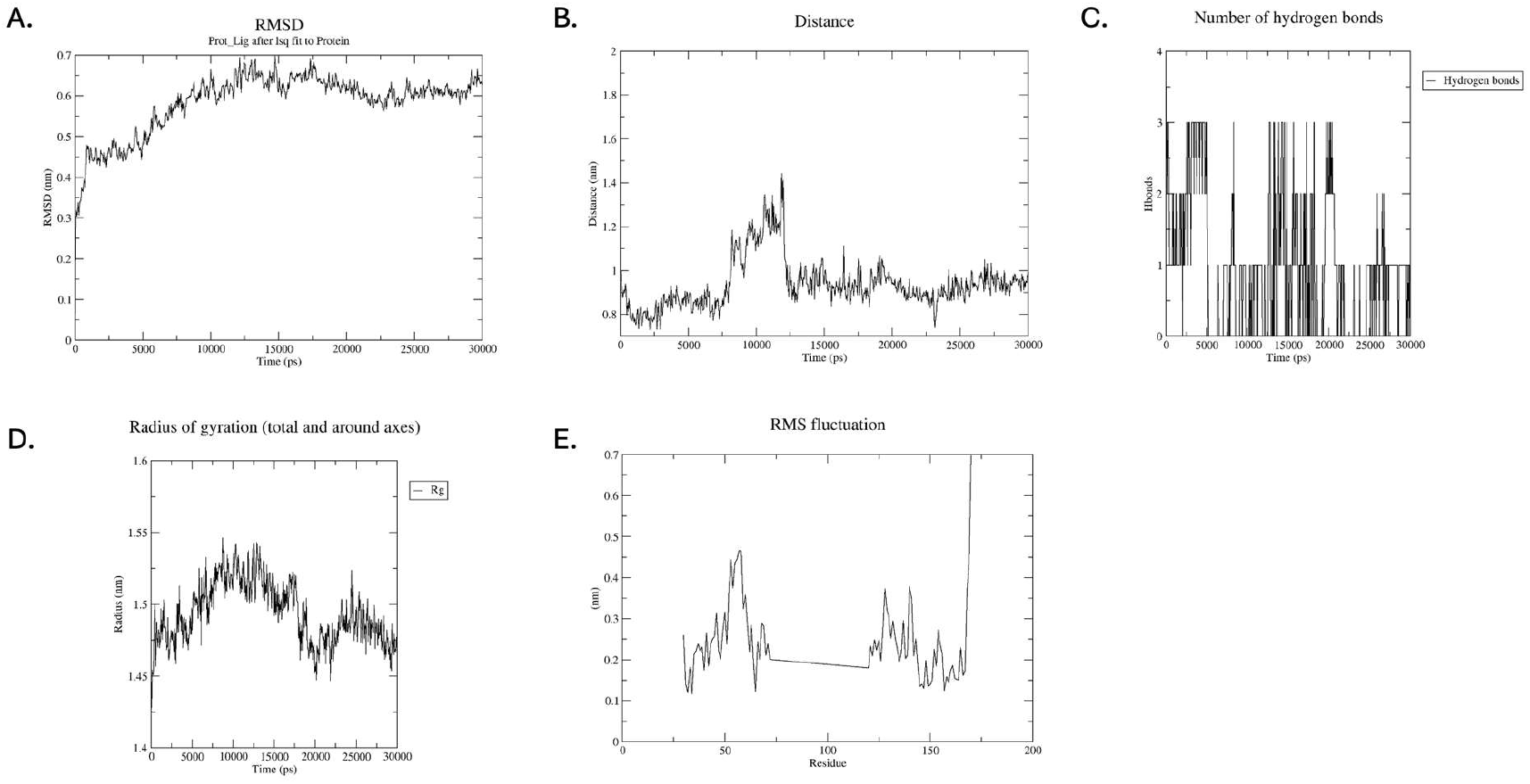
Analysis of the MD simulation of N05 bound to LAG-3 D1. **(A)** Ligand RMSD over 30 ns, showing initial adjustment followed by stabilization within the pocket. **(B)** Distance between N05 and the pocket center over time, indicating early rearrangement and subsequent stable engagement. **(C)** Number of ligand-protein hydrogen bonds throughout the trajectory, showing intermittent, dynamically formed interactions. **(D)** Radius of gyration of the protein during the simulation, indicating overall structural stability. **(E)** Residue-wise RMSF values highlighting flexible loop regions and pocket-associated motion.

Backbone RMSF analysis revealed the expected flexibility in surface-exposed loops (**Fig. 6E**), and the pocket-forming regions spanning residues ∼44-62 and ∼130-160 showed moderate mobility consistent with the transient behavior observed in the apo state. No large-scale conformational rearrangements occurred in response to ligand binding, and the radius of gyration remained stable (approximately 1.48-1.52 nm; **Fig. 6D**), supporting overall structural stability of the complex. Hydrogen-bond analysis indicated intermittent formation of zero to three hydrogen bonds between N05 and the protein (**Fig. 6C**), fluctuating over time. This behavior suggests a dynamic mixture of polar and hydrophobic interactions rather than sustained directional hydrogen bonding, consistent with the shallow, cryptic nature of the pocket and with the physicochemical profile of the ligand.

MM-GBSA analysis over the production window yielded consistently favorable total binding energies (**Fig. 7A-B**), with values predominantly between -2950 and -3100 kcal/mol. Although absolute magnitudes are not directly comparable across systems, the stable energetic profile supports the qualitative conclusion that N05 remains associated with the pocket throughout the simulation. Residue-wise decomposition highlighted contributions from residues forming the core of the cavity and stabilizing the ligand in its MD-refined pose (**Fig. 7C**), including ALA59, ARG57, GLY60, VAL61, THR62, PHE132, SER133, LEU134, GLU124, ARG127, and the C-terminal cluster composed of ALA149, ALA150, VAL151, SER159, and CYS160.

**Figure 7.**
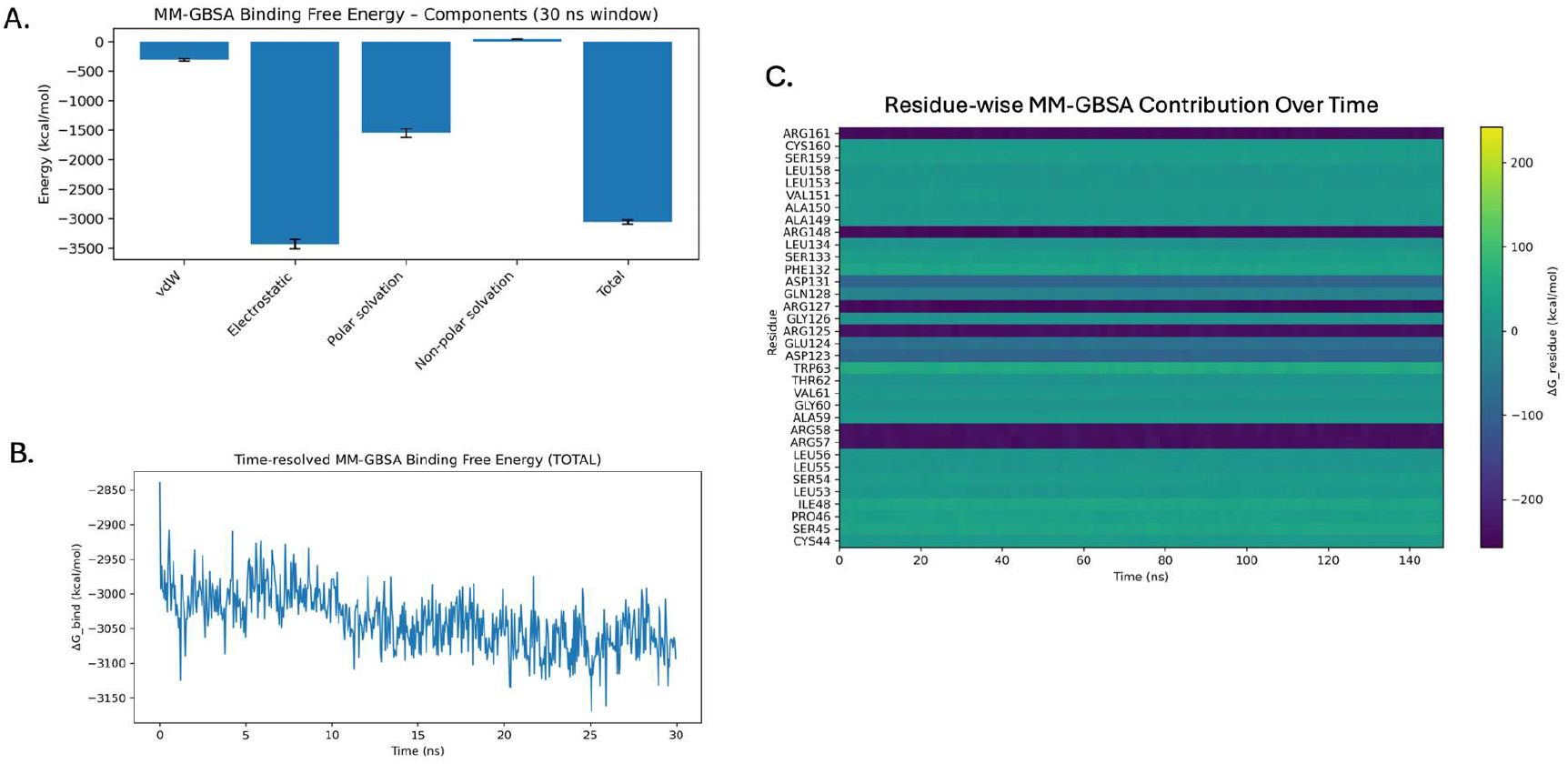
MM-GBSA analysis of N05 binding to LAG-3 D1 during MD simulation. (A) Decomposition of MM-GBSA binding free energy over the 30-ns trajectory, showing contributions from van der Waals, electrostatic, polar solvation, and nonpolar solvation terms. (B)Time-resolved total MM-GBSA binding free energy, indicating stable ligand association throughout the simulation. **(C)** Residue-wise MM-GBSA contribution heatmap highlighting residues that consistently stabilize N05 during the trajectory.

Taken together, the RMSD behavior, distance analyses, pocket dynamics, and MM-GBSA results indicate that N05 undergoes an initial pose adjustment before achieving stable engagement within the cryptic cavity revealed by the apo simulation. The ligand remains associated throughout the trajectory, supporting the feasibility of small-molecule binding at this site and complementing the biophysical and functional data identifying N05 as a tractable starting point for further optimization.

## D. Discussion

Small-molecule modulation of immune checkpoint receptors remains an underexplored but strategically important goal. In contrast to PD-1 or CTLA-4, where multiple campaigns have uncovered tractable pockets and produced early small-molecule antagonists, work on LAG-3 has centered almost entirely on monoclonal antibodies and protein-engineered modalities.^2-4^ The structural basis for this focus is apparent: the D1 domain binds MHC class II through an extended, relatively flat surface shaped by a loop insertion and adjacent β-strands,^5-9^ a configuration long considered inhospitable to classical drug-like binding. Within this context, our objective was to assess whether the LAG-3 D1 surface possesses latent ligandability that might be revealed through an MD-informed, structure-based workflow, and to determine whether small molecules can exploit conformations not evident in static structures.

Our MD simulations of isolated LAG-3 D1 showed that the domain is not rigid but samples conformations in which a shallow pocket transiently opens adjacent to the MHCII-binding face. The spontaneous appearance of this cavity during unbiased simulation indicates that the D1 surface has inherent structural plasticity capable of revealing ligand-accommodating geometries. Its recurrence across the trajectory, together with its proximity to residues involved in LAG-3/MHCII recognition,^5-8^ suggested that this region could support small-molecule engagement despite lacking a canonical cleft in available crystal structures.

Guided by this MD-revealed cavity, we applied a focused virtual screening pipeline to identify compounds capable of occupying the site. Screening ∼10,000 diverse molecules with Glide produced a small set of computational hits, from which one compound, N05, demonstrated reproducible binding in MST experiments and measurable inhibition of the LAG-3/MHCII interaction in an HTRF biochemical assay. Although its affinity lies in the micromolar range, this is typical for initial binders targeting protein-protein interfaces and aligns with early PD-1/PD-L1 small-molecule efforts.^10-14^

MD simulations of the LAG-3-N05 complex further support this conclusion. Following an initial pose adjustment, the ligand settled into the same cavity observed in the apo simulations and remained stably engaged over tens of nanoseconds. Residue-wise energy decomposition highlighted consistent stabilizing contributions from residues defining the cryptic pocket, reinforcing that the site is not a simulation artifact but a genuine structural feature capable of hosting ligand interactions with measurable biophysical and functional consequences.

Together, these findings address the central aims of this study: to determine whether LAG-3 D1 exhibits structurally accessible pockets amenable to small-molecule binding, and to evaluate whether a practical structure-guided workflow can identify compounds that exploit such pockets. Although this work does not attempt full optimization or mechanistic dissection, it provides a foundation for both. As structural and cellular studies of LAG-3 continue to expand, the demonstration that D1 can accommodate small-molecule binding opens the possibility of complementary or alternative therapeutic strategies beyond antibody blockade.

## Notes

### Competing Interest Statement

The authors have declared no competing interest.

